## Supplementary file 2 for "Varidnaviruses in the human gut: a major expansion of the order *Vinavirales*"

### Slide 1
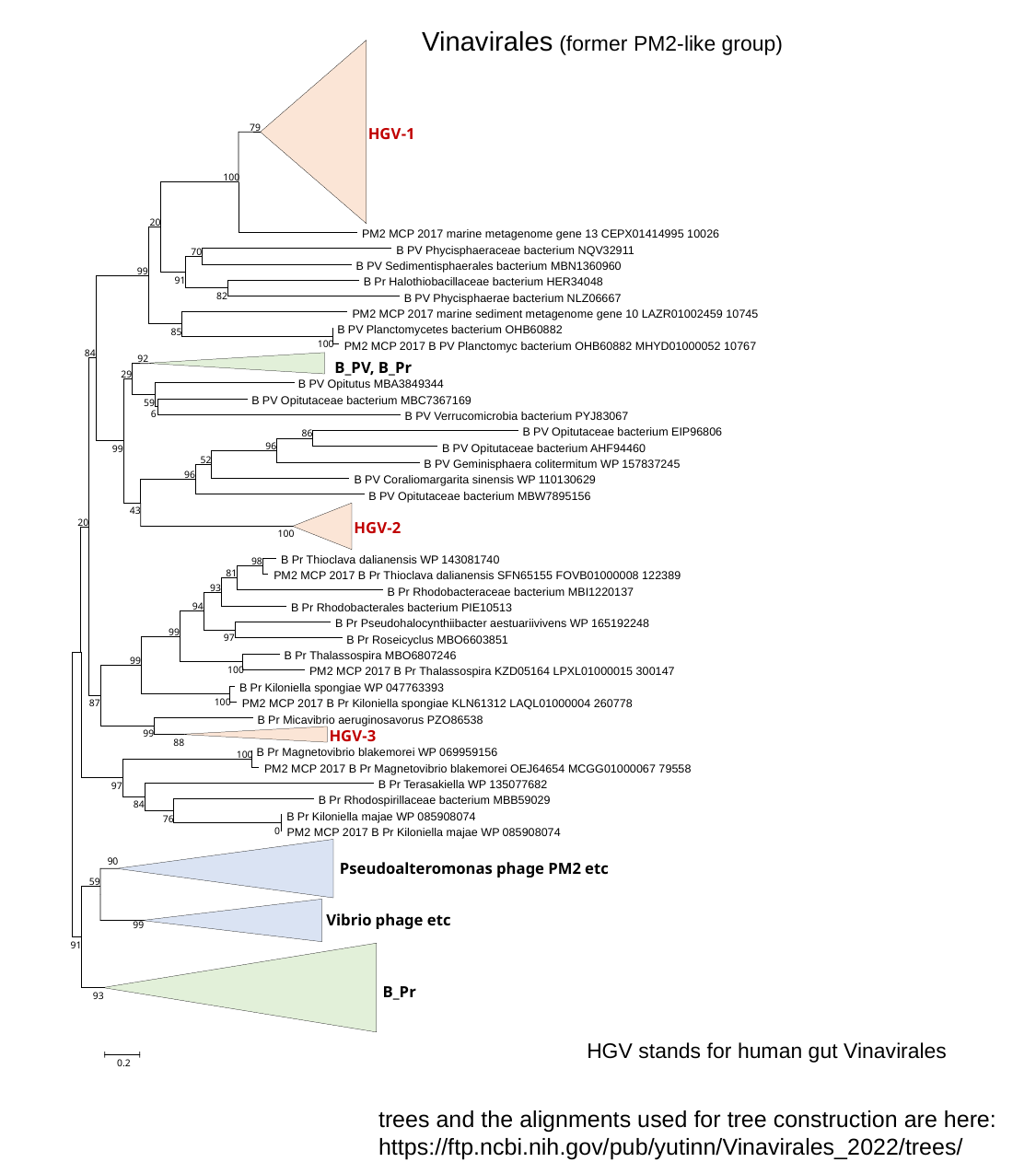

Vinavirales (former PM2-like group)
79
HGV-1
100
20
 PM2 MCP 2017 marine metagenome gene 13 CEPX01414995 10026
 B PV Phycisphaeraceae bacterium NQV32911
70
 B PV Sedimentisphaerales bacterium MBN1360960
99
91
 B Pr Halothiobacillaceae bacterium HER34048
82
 B PV Phycisphaerae bacterium NLZ06667
 PM2 MCP 2017 marine sediment metagenome gene 10 LAZR01002459 10745
 B PV Planctomycetes bacterium OHB60882
85
100
 PM2 MCP 2017 B PV Planctomyc bacterium OHB60882 MHYD01000052 10767
84
92
 B_PV, B_Pr
29
 B PV Opitutus MBA3849344
 B PV Opitutaceae bacterium MBC7367169
59
6
 B PV Verrucomicrobia bacterium PYJ83067
 B PV Opitutaceae bacterium EIP96806
86
96
 B PV Opitutaceae bacterium AHF94460
99
52
 B PV Geminisphaera colitermitum WP 157837245
96
 B PV Coraliomargarita sinensis WP 110130629
 B PV Opitutaceae bacterium MBW7895156
43
20
HGV-2
100
 B Pr Thioclava dalianensis WP 143081740
98
81
 PM2 MCP 2017 B Pr Thioclava dalianensis SFN65155 FOVB01000008 122389
93
 B Pr Rhodobacteraceae bacterium MBI1220137
94
 B Pr Rhodobacterales bacterium PIE10513
 B Pr Pseudohalocynthiibacter aestuariivivens WP 165192248
99
97
 B Pr Roseicyclus MBO6603851
 B Pr Thalassospira MBO6807246
99
100
 PM2 MCP 2017 B Pr Thalassospira KZD05164 LPXL01000015 300147
 B Pr Kiloniella spongiae WP 047763393
100
 PM2 MCP 2017 B Pr Kiloniella spongiae KLN61312 LAQL01000004 260778
87
 B Pr Micavibrio aeruginosavorus PZO86538
HGV-3
99
88
 B Pr Magnetovibrio blakemorei WP 069959156
100
 PM2 MCP 2017 B Pr Magnetovibrio blakemorei OEJ64654 MCGG01000067 79558
 B Pr Terasakiella WP 135077682
97
 B Pr Rhodospirillaceae bacterium MBB59029
84
 B Pr Kiloniella majae WP 085908074
76
0
 PM2 MCP 2017 B Pr Kiloniella majae WP 085908074
99
90
 Pseudoalteromonas phage PM2 etc
59
 Vibrio phage etc
91
 B_Pr
93
0.2
HGV stands for human gut Vinavirales
trees and the alignments used for tree construction are here:
https://ftp.ncbi.nih.gov/pub/yutinn/Vinavirales_2022/trees/

### Slide 2
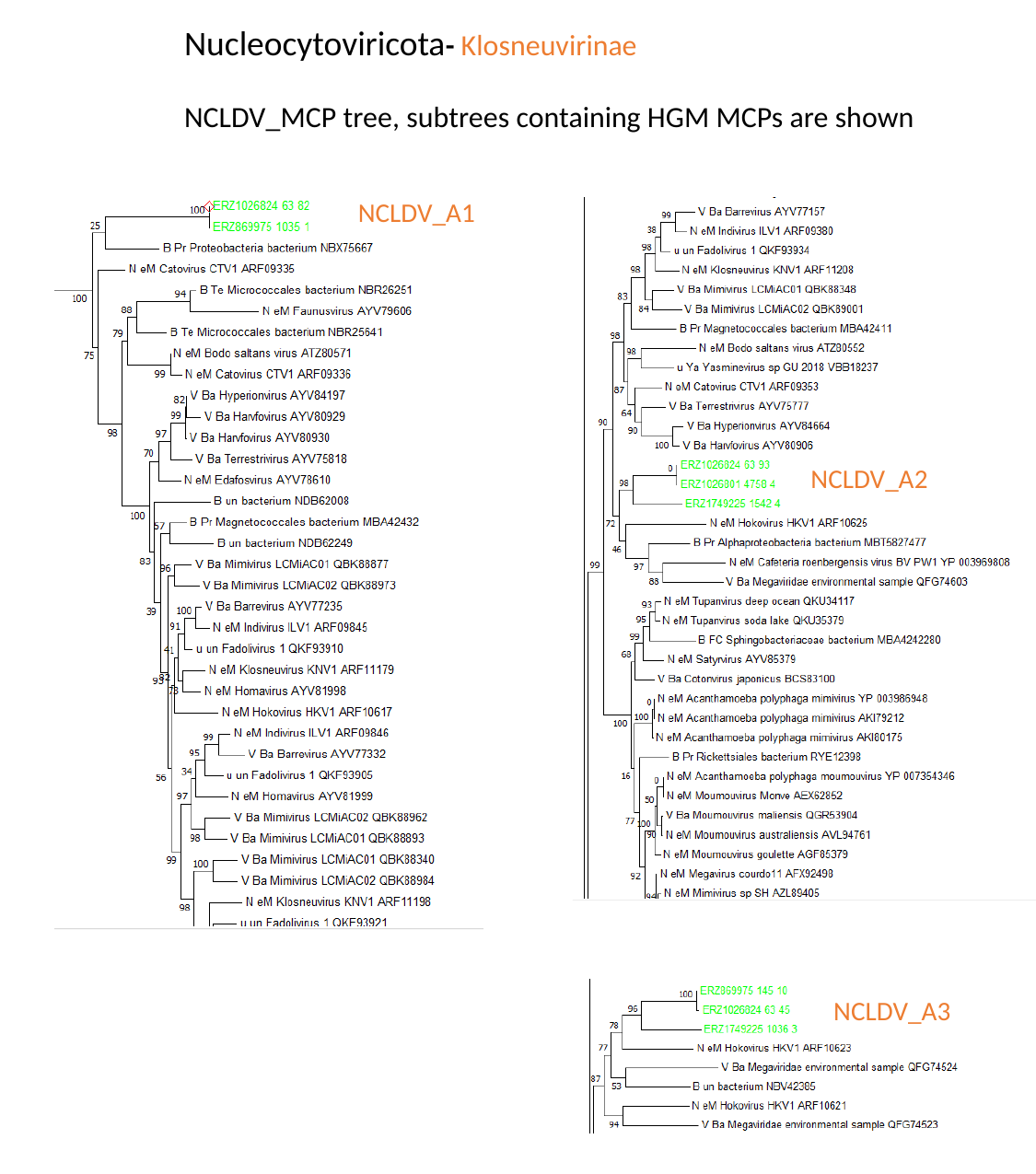

Nucleocytoviricota- Klosneuvirinae
NCLDV_MCP tree, subtrees containing HGM MCPs are shown
NCLDV_A1
NCLDV_A2
NCLDV_A3

### Slide 3
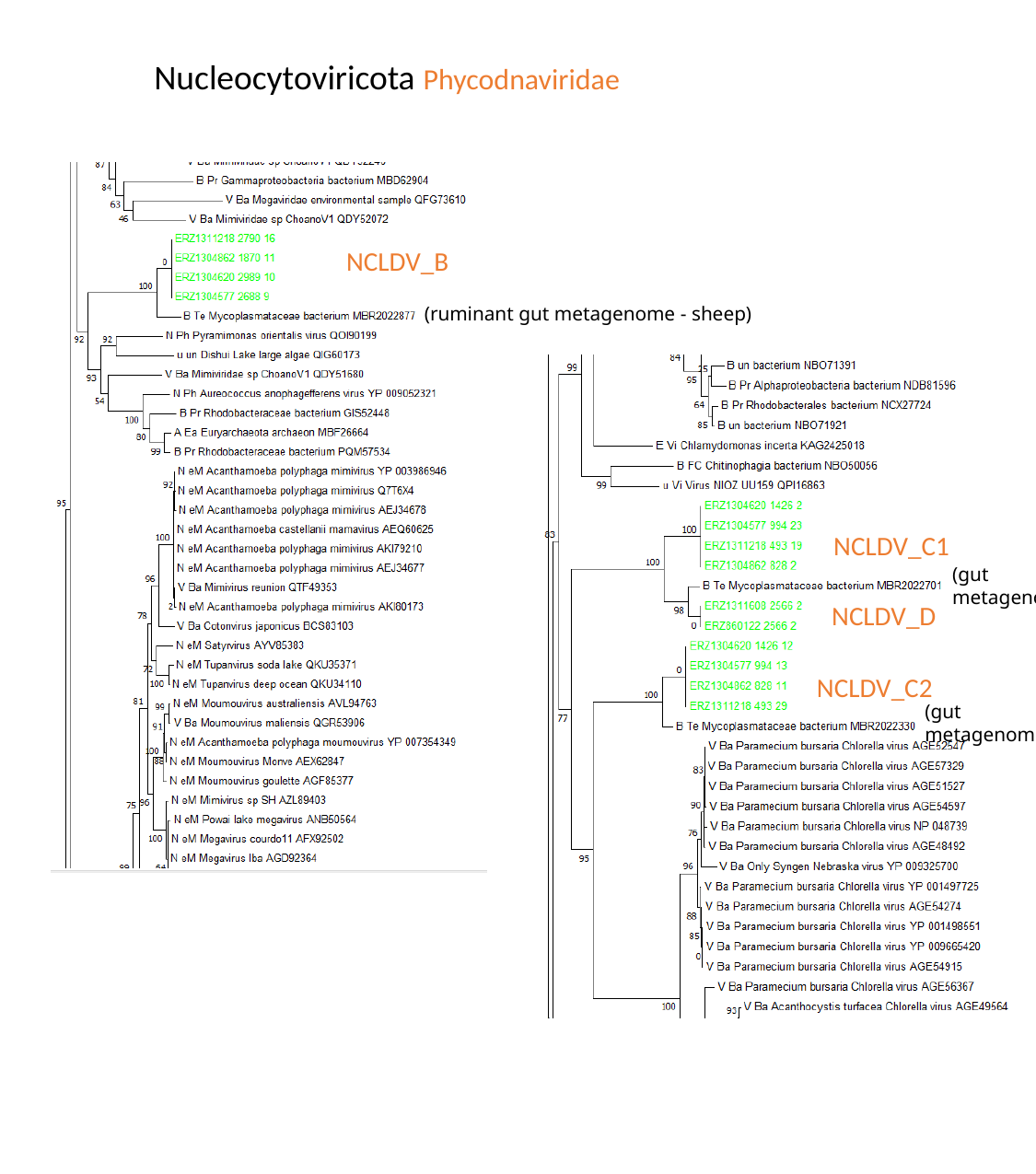

Nucleocytoviricota Phycodnaviridae
NCLDV_B
(ruminant gut metagenome - sheep)
NCLDV_C1
NCLDV_D
NCLDV_C2
(gut metagenome)
(gut metagenome)

### Slide 4
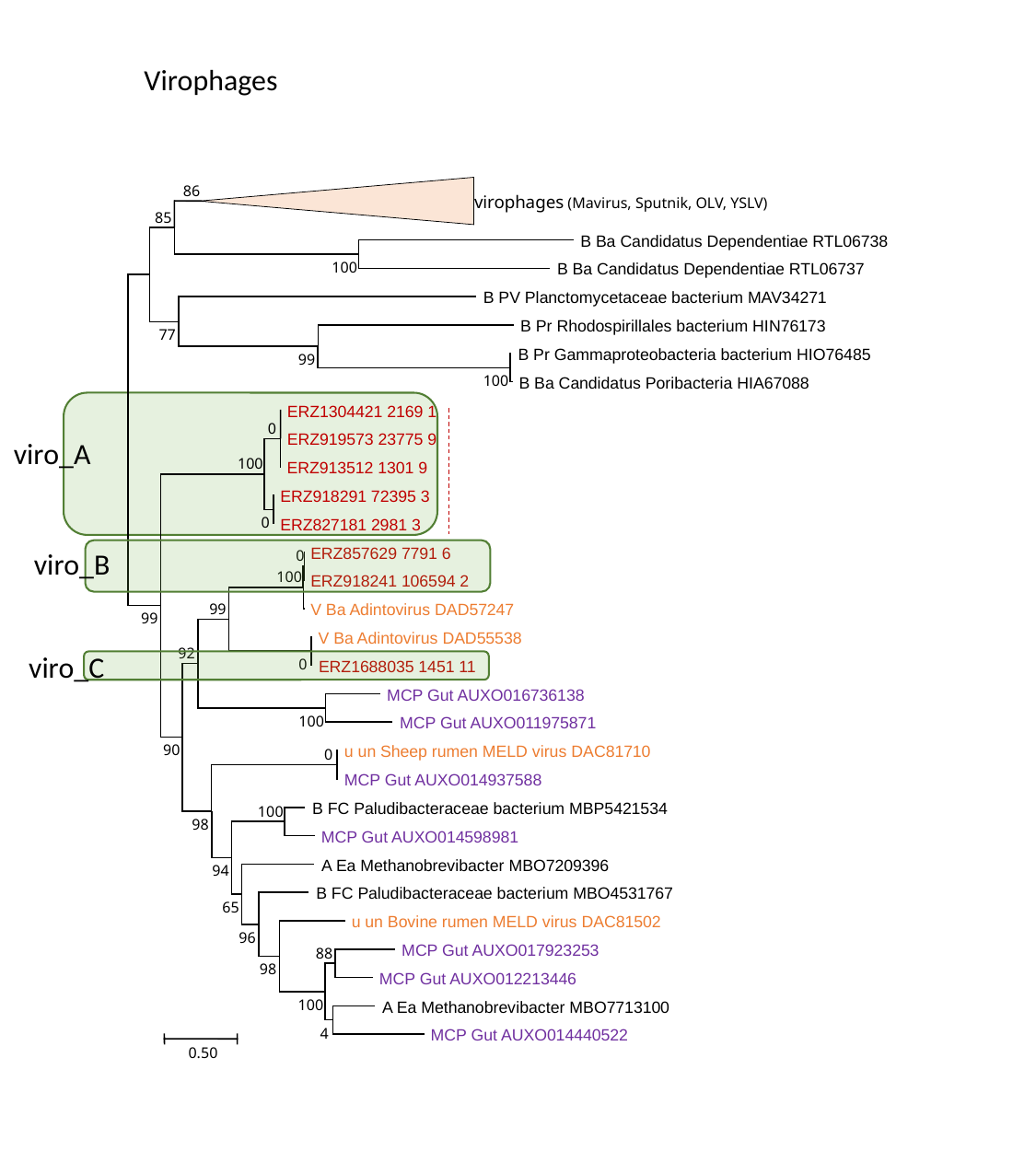

Virophages
86
 virophages (Mavirus, Sputnik, OLV, YSLV)
85
 B Ba Candidatus Dependentiae RTL06738
100
 B Ba Candidatus Dependentiae RTL06737
 B PV Planctomycetaceae bacterium MAV34271
 B Pr Rhodospirillales bacterium HIN76173
77
 B Pr Gammaproteobacteria bacterium HIO76485
99
100
 B Ba Candidatus Poribacteria HIA67088
 ERZ1304421 2169 1
0
 ERZ919573 23775 9
100
 ERZ913512 1301 9
 ERZ918291 72395 3
0
 ERZ827181 2981 3
 ERZ857629 7791 6
0
100
 ERZ918241 106594 2
 V Ba Adintovirus DAD57247
99
99
 V Ba Adintovirus DAD55538
92
0
 ERZ1688035 1451 11
 MCP Gut AUXO016736138
100
 MCP Gut AUXO011975871
90
 u un Sheep rumen MELD virus DAC81710
0
 MCP Gut AUXO014937588
 B FC Paludibacteraceae bacterium MBP5421534
100
98
 MCP Gut AUXO014598981
 A Ea Methanobrevibacter MBO7209396
94
 B FC Paludibacteraceae bacterium MBO4531767
65
 u un Bovine rumen MELD virus DAC81502
96
 MCP Gut AUXO017923253
88
98
 MCP Gut AUXO012213446
100
 A Ea Methanobrevibacter MBO7713100
4
 MCP Gut AUXO014440522
0.50
viro_A
viro_B
viro_C

### Slide 5
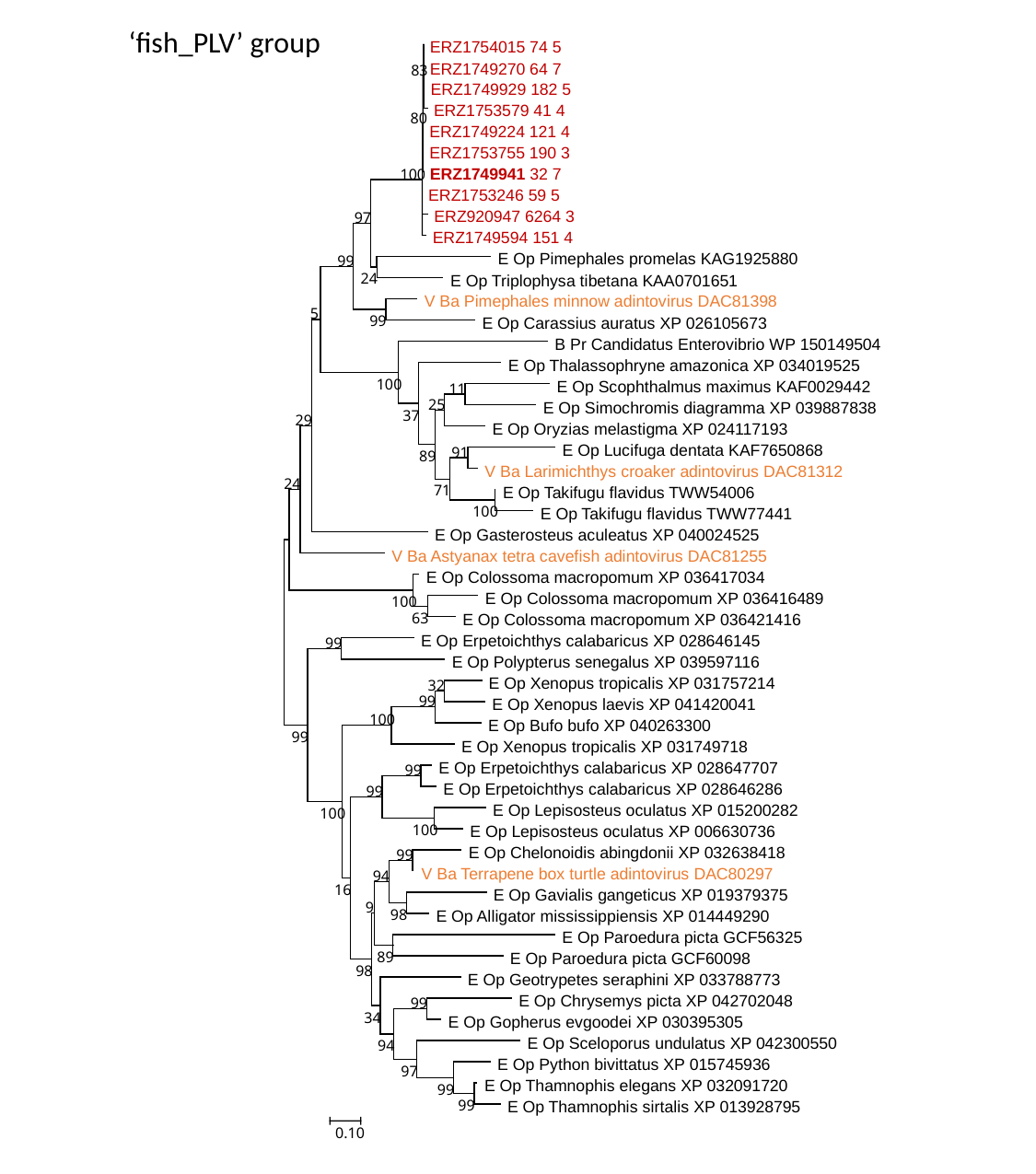

‘fish_PLV’ group
 ERZ1754015 74 5
 ERZ1749270 64 7
83
 ERZ1749929 182 5
 ERZ1753579 41 4
80
 ERZ1749224 121 4
 ERZ1753755 190 3
 ERZ1749941 32 7
100
 ERZ1753246 59 5
 ERZ920947 6264 3
97
 ERZ1749594 151 4
 E Op Pimephales promelas KAG1925880
99
24
 E Op Triplophysa tibetana KAA0701651
 V Ba Pimephales minnow adintovirus DAC81398
5
99
 E Op Carassius auratus XP 026105673
 B Pr Candidatus Enterovibrio WP 150149504
 E Op Thalassophryne amazonica XP 034019525
100
 E Op Scophthalmus maximus KAF0029442
11
25
 E Op Simochromis diagramma XP 039887838
37
29
 E Op Oryzias melastigma XP 024117193
 E Op Lucifuga dentata KAF7650868
91
89
 V Ba Larimichthys croaker adintovirus DAC81312
24
71
 E Op Takifugu flavidus TWW54006
100
 E Op Takifugu flavidus TWW77441
 E Op Gasterosteus aculeatus XP 040024525
 V Ba Astyanax tetra cavefish adintovirus DAC81255
 E Op Colossoma macropomum XP 036417034
 E Op Colossoma macropomum XP 036416489
100
63
 E Op Colossoma macropomum XP 036421416
 E Op Erpetoichthys calabaricus XP 028646145
99
 E Op Polypterus senegalus XP 039597116
 E Op Xenopus tropicalis XP 031757214
32
99
 E Op Xenopus laevis XP 041420041
100
 E Op Bufo bufo XP 040263300
99
 E Op Xenopus tropicalis XP 031749718
 E Op Erpetoichthys calabaricus XP 028647707
99
 E Op Erpetoichthys calabaricus XP 028646286
99
 E Op Lepisosteus oculatus XP 015200282
100
100
 E Op Lepisosteus oculatus XP 006630736
 E Op Chelonoidis abingdonii XP 032638418
99
 V Ba Terrapene box turtle adintovirus DAC80297
94
16
 E Op Gavialis gangeticus XP 019379375
9
98
 E Op Alligator mississippiensis XP 014449290
 E Op Paroedura picta GCF56325
89
 E Op Paroedura picta GCF60098
98
 E Op Geotrypetes seraphini XP 033788773
 E Op Chrysemys picta XP 042702048
99
34
 E Op Gopherus evgoodei XP 030395305
 E Op Sceloporus undulatus XP 042300550
94
 E Op Python bivittatus XP 015745936
97
 E Op Thamnophis elegans XP 032091720
99
99
 E Op Thamnophis sirtalis XP 013928795
0.10

### Slide 6
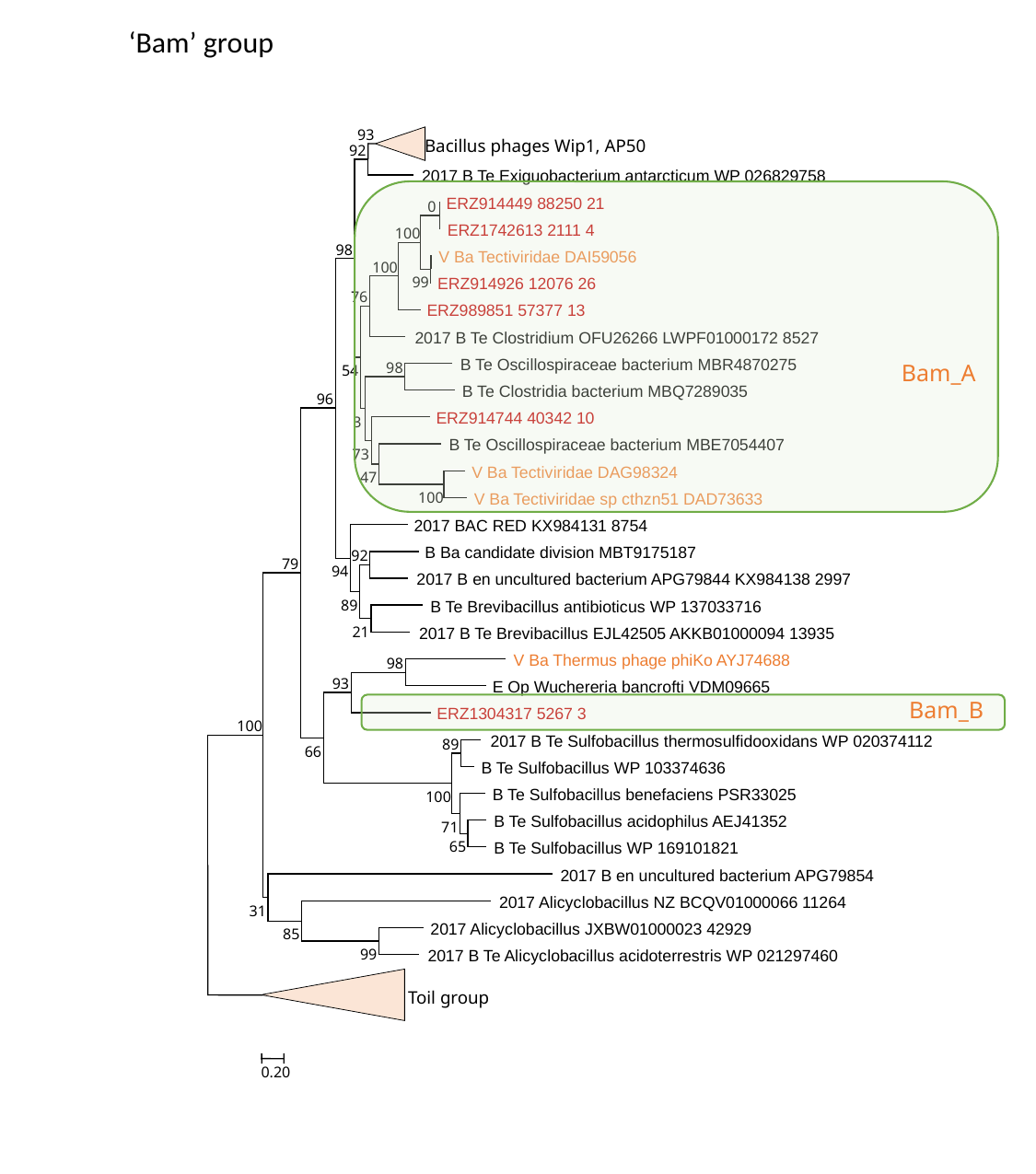

‘Bam’ group
93
 Bacillus phages Wip1, AP50
92
 2017 B Te Exiguobacterium antarcticum WP 026829758
 ERZ914449 88250 21
0
 ERZ1742613 2111 4
100
98
 V Ba Tectiviridae DAI59056
100
99
 ERZ914926 12076 26
76
 ERZ989851 57377 13
 2017 B Te Clostridium OFU26266 LWPF01000172 8527
 B Te Oscillospiraceae bacterium MBR4870275
98
54
 B Te Clostridia bacterium MBQ7289035
96
 ERZ914744 40342 10
3
 B Te Oscillospiraceae bacterium MBE7054407
73
 V Ba Tectiviridae DAG98324
47
100
 V Ba Tectiviridae sp cthzn51 DAD73633
 2017 BAC RED KX984131 8754
 B Ba candidate division MBT9175187
92
79
94
 2017 B en uncultured bacterium APG79844 KX984138 2997
89
 B Te Brevibacillus antibioticus WP 137033716
21
 2017 B Te Brevibacillus EJL42505 AKKB01000094 13935
 V Ba Thermus phage phiKo AYJ74688
98
93
 E Op Wuchereria bancrofti VDM09665
 ERZ1304317 5267 3
100
 2017 B Te Sulfobacillus thermosulfidooxidans WP 020374112
89
66
 B Te Sulfobacillus WP 103374636
 B Te Sulfobacillus benefaciens PSR33025
100
 B Te Sulfobacillus acidophilus AEJ41352
71
65
 B Te Sulfobacillus WP 169101821
 2017 B en uncultured bacterium APG79854
 2017 Alicyclobacillus NZ BCQV01000066 11264
31
 2017 Alicyclobacillus JXBW01000023 42929
85
99
 2017 B Te Alicyclobacillus acidoterrestris WP 021297460
 Toil group
0.20
Bam_A
Bam_B

### Slide 7
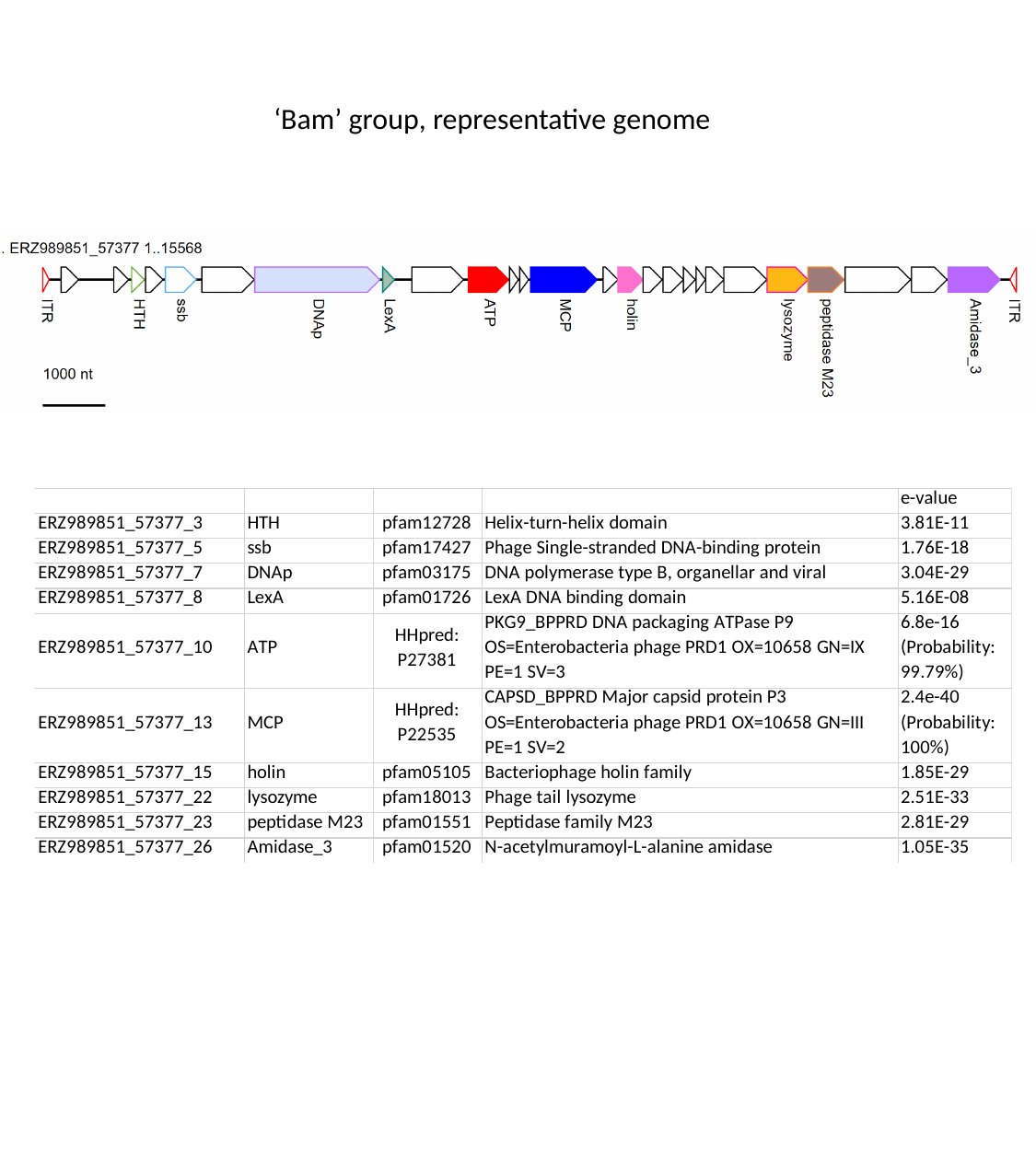

‘Bam’ group, representative genome
