## Supplementary file 8 for "Varidnaviruses in the human gut: a major expansion of the order *Vinavirales*"

### Slide 1
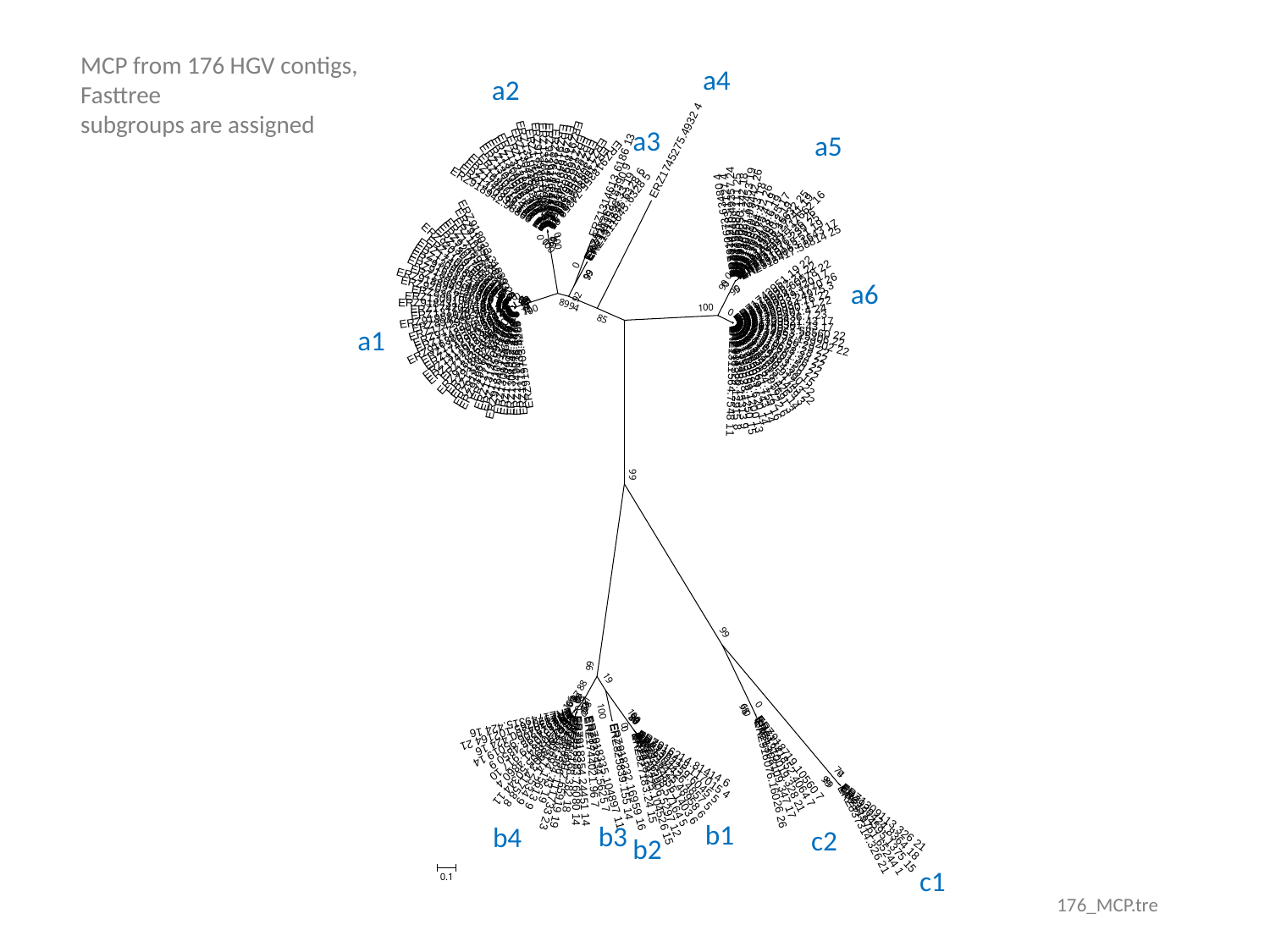

MCP from 176 HGV contigs, Fasttree
subgroups are assigned
a4
a2
a3
a5
a6
a1
b1
b3
b4
c2
b2
c1
176_MCP.tre

### Slide 2
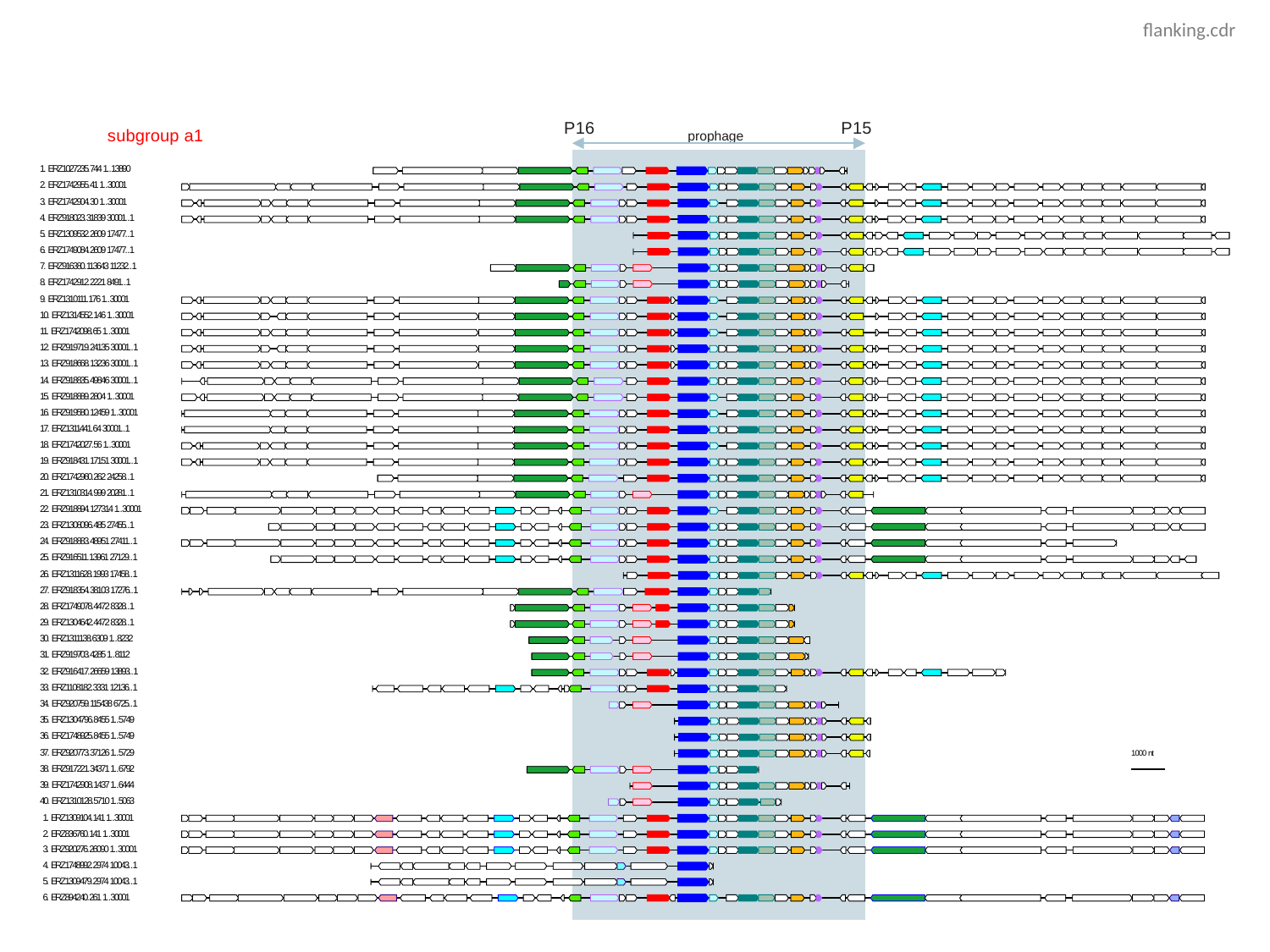

flanking.cdr

### Slide 3
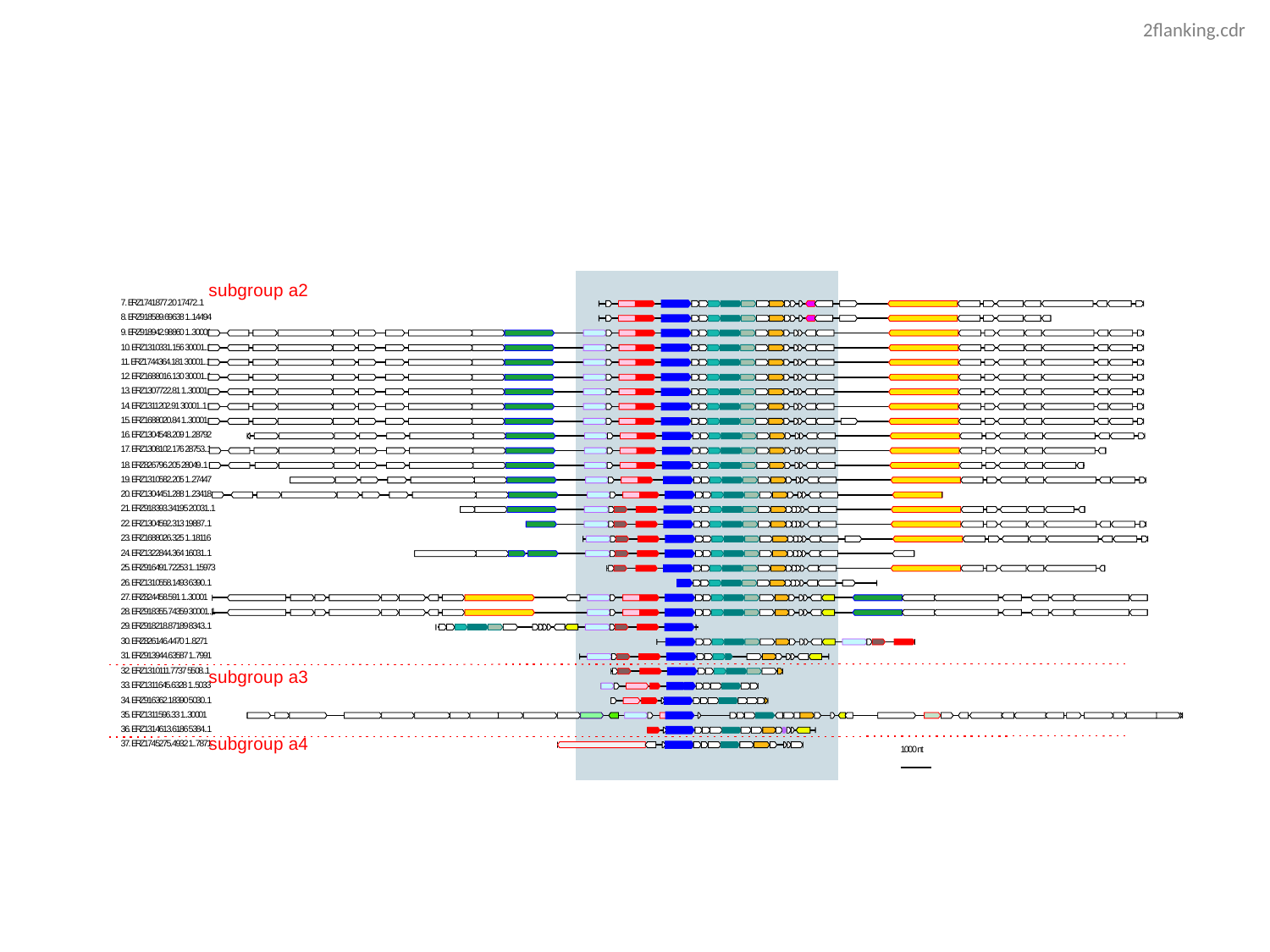

2flanking.cdr

### Slide 4
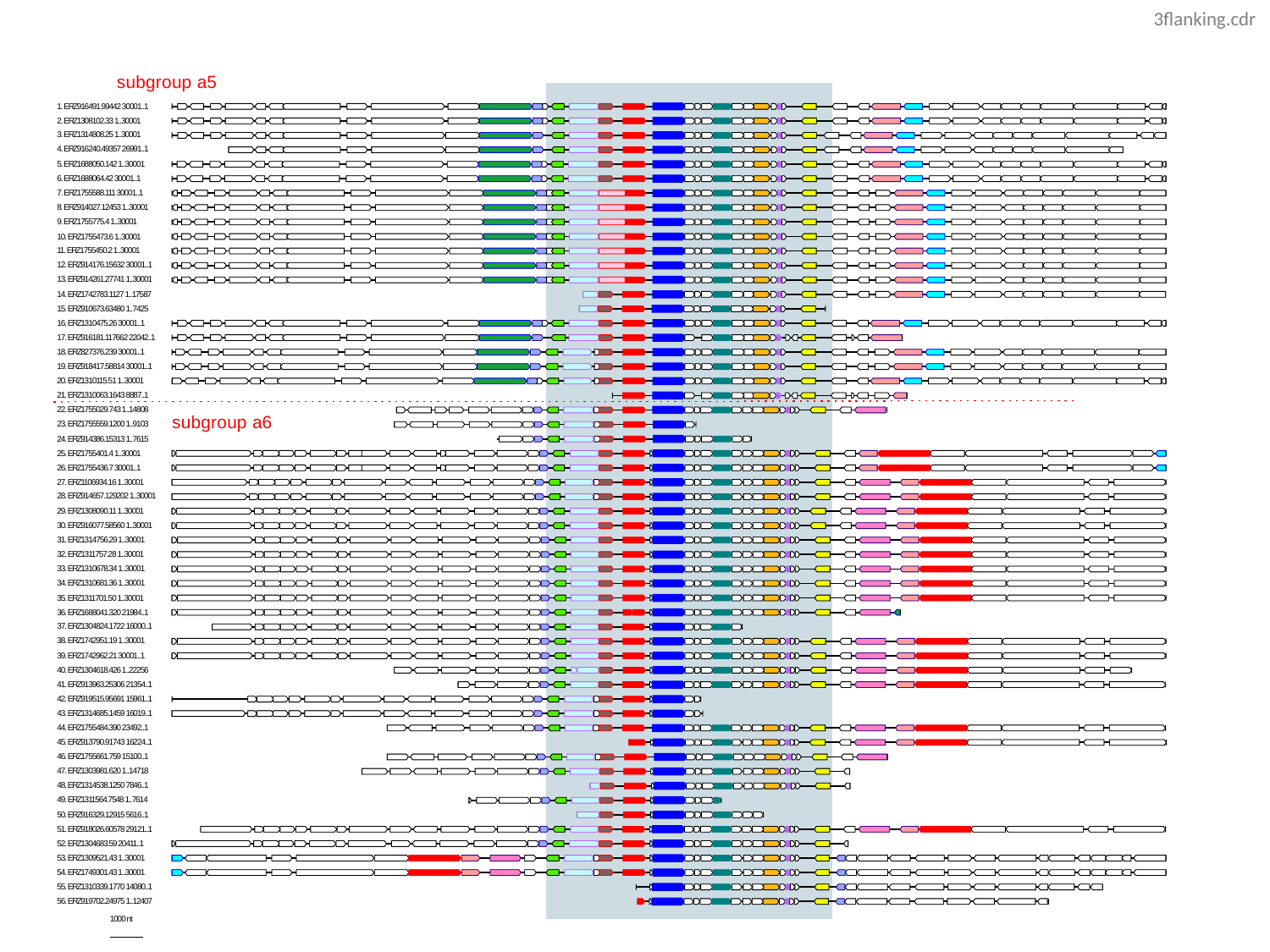

3flanking.cdr

### Slide 5
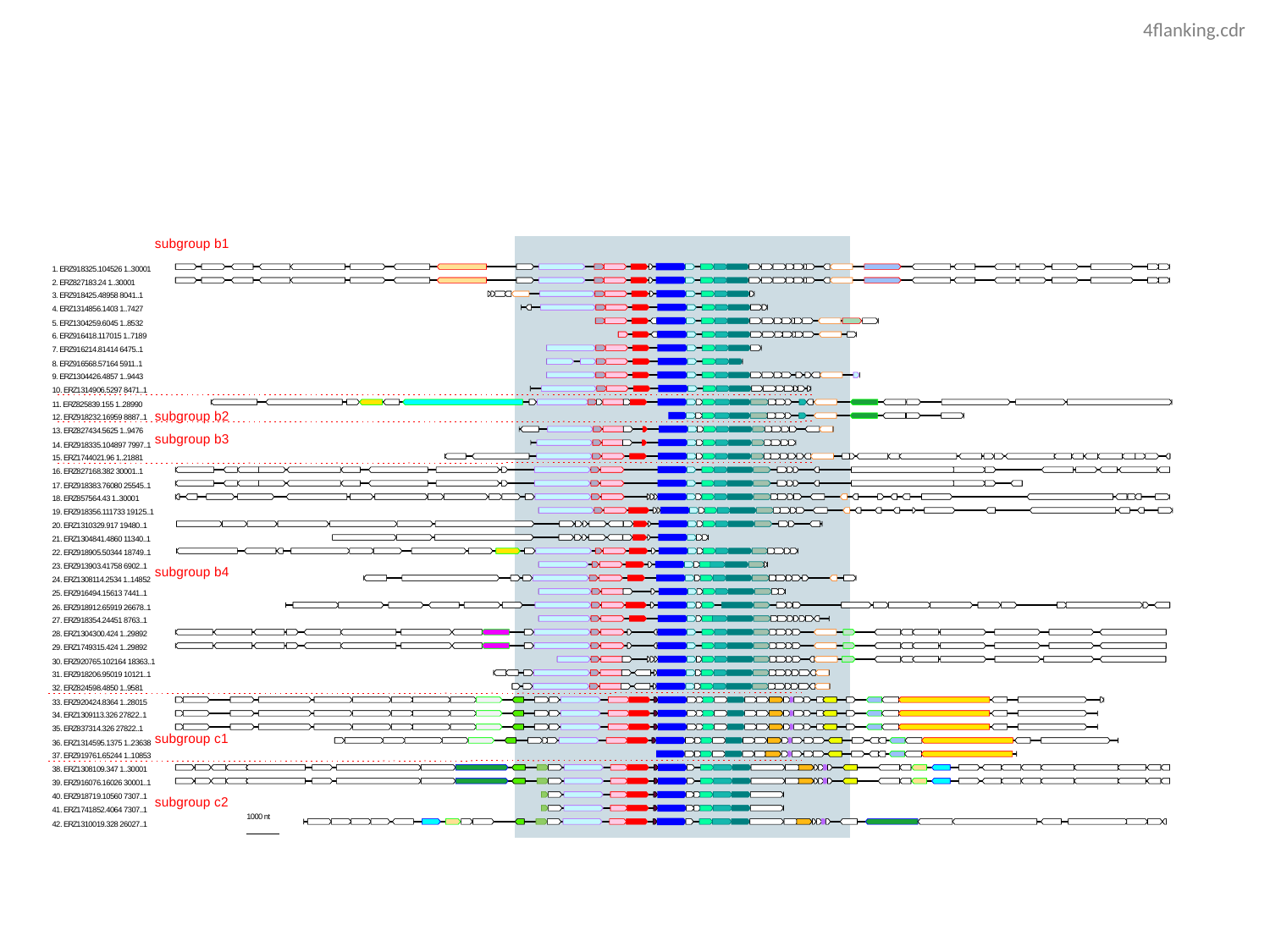

4flanking.cdr
